# Measuring Mitochondrial Electron Transfer Complexes in Previously Frozen Cardiac Tissue from the Offspring of Sow: A Model to Assess Exercise-Induced Mitochondrial Bioenergetics Changes

**DOI:** 10.1101/2021.06.08.447578

**Authors:** Daniel Barrera, Sierra Upton, Megan Rauch, Tara Notarianni, Ki Suk Eum, Megan Liberty, Sarmila Venkoba Sah, Robert Liu, Sean Newcomer, Linda E. May, Emre Agbas, Jessica Sage, Edina Kosa, Abdulbaki Agbas

**Affiliations:** AdventHelath, Orlando, FL, United States; University of Illinois Chicago School of Public Health, Illinois, United States; Lincoln Memorial University, Harrogate, TN, United States; Children’s Mercy Hospital, Kansas City, MO, United States; Scripps Clinic, San Diego, CA, United States; Cincinnati Children’s Hospital Medical Center, Ohio, United States; Kansas City University, Kansas City, MO, United States; Roblex Rex Veterans Affairs Medical Center, Louisville, KY, United states; California State University San Marcos, CA, United States; East Carolina University, Greenville, NC, United States; Massachusetts Institute of Technology, Cambridge, MA, United States; Boehringer Ingelheim Norway KS, St. Joseph, MO, United States; Heartland Center for Mitochondrial Medicine, Kansas City, KS United States

## Abstract

Measuring the mitochondrial electron transfer complex (ETC) profile from previously frozen heart tissue samples from offspring born to an exercised sow provided descriptive data about exercise-induced mitochondrial biochemical changes in heart tissue from the offspring born to the exercised sow. The hypothesis proposed and tested was that a regular maternal exercise of a sow during pregnancy would increase the mitochondrial efficiency of offspring heart bioenergetics. This hypothesis was tested by isolating mitochondria using a mild-isolation procedure to assess mitochondrial ETC and supercomplex profiles. The procedure described here allowed for the processing of previously frozen archived heart tissues and eliminated the necessity of fresh mitochondria preparation for the assessment of mitochondrial ETC complexes, supercomplexes, and ETC complex activity profiles. This protocol describes the optimal ETC protein complex measurement in multiplexed antibody-based immunoblotting and super complex assessment using blue-native gel electrophoresis.

**SUMMARY:** Preparation of mitochondria-enriched samples from previously frozen archived solid tissues allowed the investigators to perform both functional and analytical assessments of mitochondria in various experimental modalities. This study demonstrates how to prepare mitochondria-enriched preparations from frozen heart tissue and perform analytical assessments of mitochondria.

## INTRODUCTION

The goal of this protocol was to provide detailed steps to obtain mitochondria-enriched preparation from previously frozen heart tissue with a new technology of low energy mechanical disruption of tissue that improves tissue lysis and extraction of mitochondria. With this method, improved extraction efficiency without generating high shear stress or high temperature and short homogenization time (10–12 s) become achievable^1^.

To obtain mitochondria from archived frozen tissue is a valuable asset to perform both functional^2^ and biochemical studies^3^ otherwise not easily repeatable under the exact experimental conditions. A classic Potter-Elvehjem Teflon pestle glass homogenizer or Dounce homogenizer has been used and is still being used in research laboratories to homogenize soft tissues such as liver, kidney, and brain. However, homogenizing hard tissues such as muscle and heart require more homogenization time, enzyme treatment, high-speed homogenization, and extensive user experience. This is disadvantageous for extracting intact organelles such as mitochondria from hard tissue such as muscle and heart. The method described in this protocol is used to obtain high yield mitochondria-enriched preparation to analyze mitochondrial electron transport chain (ETC) protein complexes and their supercomplex formation in heart tissues harvested from offspring born to exercised and sedentary sow, flash-frozen in liquid nitrogen, and stored at −80 °C for future use. This method allows the user to isolate mitochondria enriched preparation from previously frozen archived tissues.

External nanomaterial exposure to pregnant rodents can negatively affect cardiac function and mitochondrial respiration and bioenergetics on progeny during gestation^4^. Nevertheless, aerobic exercise-induced positive changes in fetal myocyte bioenergetics during pregnancy are yet to be documented. However, emerging studies provide evidence that maternal aerobic exercise during pregnancy has a positive influence on fetal cardiac function^5^. In order to provide further evidence, an analysis of the longitudinal effects of maternal exercise on offspring cardiac mitochondrial respiratory chain complexes (i.e., Complex I through Complex V) during pregnancy was performed.

This study has significant health relevance since the results may suggest that maternal exercise improves the efficiency of energy production in the cardiac mitochondria of the offspring. In this study, sows (female pig) were used as an animal model for two reasons: (i) cardiac physiology is similar to human^6^, and (ii) heart tissue harvest from offspring from different time points is feasible under an institutional IACUC approval. The proposed study aims to answer many of the fundamental questions linking maternal exercise and its potential positive effects on the cellular and biochemical makeup of the offspring’s heart tissue. This approach requires gentle yet effective mitochondria isolation techniques from previously frozen cardiac tissue obtained from the lengthy and costly longitudinal studies that addressed the issues of the bioenergetic changes within fetal cardiac myocytes during the pregnancy. The method described in this study allows utilizing large sums of previously frozen archived tissue for mitochondria-enriched preparation for both analytical and functional studies. The study will also help fill the knowledge gap in this field by providing preliminary data, which could lead to future studies determining the effects of maternal exercise on heart health in utero and beyond.

## PROTOCOL

Frozen offspring heart tissues were received from Dr. Sean Newcomer along with the institutional IACUC approval letter. The heart tissues were obtained from a long-term longitudinal study, flash-frozen in liquid nitrogen, and stored at −80 °C for future use. All protocols concerning the processing of offspring heart tissue followed the guidelines of Kansas City University IBC and IACUC committees.

### 1. Preparation of buffers and reagents

NOTE: Prepare all samples as per the manufacturer’s guidelines. Use ultra-purified water or equivalent in all recipes and protocol steps. Wear personal protection equipment (lab gloats, facemask, gloves, and goggles/face shield) during this procedure. Buffer volumes are suitable for processing six tissue samples.

1.1. Prepare 1.5 L tissue washing buffer by combining 154.035 g of sucrose (0.3 M final), 3.904 g of HEPES (10 mM final), 0.111 g of EDTA (0.2 mM final). Add ingredients to 1.0 L of water while on the stir plate, adjust the pH to 7.2 with dilute HCl, and makeup to final volume to 1.5 L. NOTE: Approximately 200 mL of washing buffer is needed per sample (50–300 mg). Six tissue samples can be processed in a day. Use the remaining washing buffer to make an isolation buffer.
1.2. Prepare the isolation buffer from the wash buffer previously described in step 1.1 by adding 210 mg of BSA in 210 mL of wash buffer. Adjust the pH to 7.4 with dilute NaOH solution. NOTE: Approximately 35 mL of isolation buffer is needed per tissue sample; therefore, 210 mL of isolation buffer is required for six samples. Prepare this buffer by adding fresh BSA solution (1mg/mL final) on the day of preparation. BSA is used to isolate mitochondria as it improves the mitochondrial properties by removing natural uncouplers such as free fatty acids. BSA has been found to have antioxidant activity by increasing oxidation rate of substrates, especially succinate, in brain mitochondria^7^. Alternatively, bovine serum albumin can be substituted by less expensive acid-free analogs such as ovalbumin.
1.3. Prepare 150 mL of resuspension buffer by combining 0.011 g of EDTA (final concentration of 0.2 mM EDTA or use 150 μL of 200 mM stock EDTA solution for 150 mL of resuspension buffer), 12.836 g of sucrose (0.25 M sucrose final concentration), and 0.236 g of Tris (10 mM Tris-HCl final). Adjust the pH to 7.8 with 0.1 N NaOH. Add 60 μL of the protease inhibitor cocktail just before use.
1.4. Prepare the trypsin solution by dissolving 7.5 mg of trypsin in 3.0 mL of 1 mM HCI (2.5 mg of trypsin/mL final). This amount is sufficient for one sample. NOTE: Prepare the trypsin solution based on the number of tissue samples planned to be processed.
1.5. Prepare the trypsin inhibitor solution from soybean by combining 13.0 mg of trypsin inhibitor in 20 mL of isolation buffer containing BSA (0.65 mg/mL of trypsin inhibitor final). NOTE: Prepare a fresh trypsin inhibitor solution.
1.6. Prepare the ammonium persulfate reagent by adding 0.1 g of ammonium persulfate in 1 mL of water (10% ammonium persulfate final). NOTE: The ammonium persulfate reagent should be either freshly prepared or a large volume can be aliquoted and kept in a −20 °C freezer.
1.7. Prepare the aminocaproic acid solution (ACA) by combining 0.131 g of 6-amino caproic acid in 1.0 mL of ultra-purified water and store it in 4 °C (1 M 6-aminocaproic acid final).
1.8. Prepare the Coomassie brilliant blue dye solution by dissolving 1 g of Coomassie G-250 in 0.5 mL of water, and 0.5 mL of 1 M aminocaproic acid (10% dye solution final concentration).
1.9. Prepare 5% Digitonin for BN-PAGE. Approximately calculate how much digitonin is needed in the sample preparation plus some extra volume to offset the pipetting errors (overestimate by 10 mg). For 15 mg: First, dilute 15 mg of digitonin in 30 μL of 100% DMSO. Vortex to help the digitonin to get into the solution. Next, add the remaining 90% of ddH_2_O (270 μL). Gently heat to mix the solution (vortex), and then cool it on ice.
1.10. Prepare the native PAGE anode buffer. Add 50 mL of 20x native PAGE running buffer to 950 mL water to make a final volume of 1 L. Prepare fresh anode buffer for immediate use. Use this as a 1x blue cathode buffer.
1.11. Prepare a dark blue cathode buffer. Dissolve 0.0160 g of Coomassie brilliant blue G-250 in 80 mL of native PAGE anode buffer; mix well. NOTE: Prepare fresh cathode buffer for immediate use. Only 60 mL may be needed but make an extra 20 mL in the event of any leaks.
1.12. Prepare the light blue cathode buffer. Add 20 mL of dark blue cathode buffer into 180 mL of native PAGE anode buffer and mix well. NOTE: This buffer can only be used once.
1.13. Prepare 5% Coomassie G-250 as a sample additive. Dilute 200 μL of 10% Coomassie stock already made (step 1.8) with 200 μL of 1 M ACA and store at the 4 °C.
1.14. Prepare 4x native PAGE sample buffer by combining 40% glycerol, 0.04% Ponceau S, 25 mM Tris-HCl (pH 6.8), 192 mM glycine in 10 mL of final volume. Aliquot them and store at −20 °C in the freezer.

### 2. Mitochondria isolation from frozen heart tissue

NOTE: Perform the mitochondria isolation procedure in a 4 °C cold room. However, in case of unavailability of the cold room, use a large size ice bath to perform the procedure.

2.1. Take out previously frozen heart tissue samples from the −80 °C freezer. NOTE: A maximum of four tissue samples may be comfortably processed for mitochondria isolation in a given day.
2.2. Weigh the tube containing the heart tissue sample and record the weight. Prepare three beakers (each 50 mL) containing 40 mL of washing buffer. Cool each beaker in the ice-salt bath with equal amounts of ice and salt mixed for 3–5 min before proceeding to the next step. Put magnetic stirring bars in each beaker and set the stirrer for medium speed. NOTE: Do not over-freeze the beakers and use them immediately just after clouds of ice crystals start to appear.
2.3. Put the frozen heart tissue directly into the ice-cold solution in the first beaker. Wait approximately 5 min while stirring and applying slight pressure to the tissue using tweezers to get out as much blood as possible.
2.4. Weigh the empty tube and subtract from the original weight (step 2.2) to obtain a net weight of the heart tissue.
2.5. Take the tissue out with forceps and squeeze the heart with a clean paper towel of filter paper to remove blood. Then, place the tissue into the second beaker. Wait approximately 5 min while stirring to remove excess blood.
2.6. Gently dry the tissue on a paper towel. Remove all fat, clotted blood, auricles, and fasciae (white tissue). Then, pool the ventricular tissue.
2.7. Shred the tissue samples using the shredder by following steps 2.7.1–2.7.9.

2.7.1. Place the heart tissue (~50–300 mg) into the tissue shredder chamber from the ram side.
2.7.2. Add ~0.2–0.8 mL of washing buffer to the cap side of the tube. NOTE: Ensure that the total volume of the sample and the washing buffer combined during the shredding does not exceed 0.7–0.8 mL.
2.7.3. Use the tube tool to cap the shredder chamber with the shredder cap snugly. NOTE: Do not over tighten the shredder cap.
2.7.4. Place the shredder chamber into the shredder holder unit with the ram side down. The ram will engage a fitting in the shredder holder.
2.7.5. Insert the bit of the shredder driver; let the bit engage with the shredder cap. Make sure that the bit is snugly locked into place. NOTE: Always use the fully charged shredder driver.
2.7.6. Manually apply pressure downward to the shredder driver and shredder chamber until the pressure indicator on the shredder holder reaches a setting of approximately 2 on the shredder holder. NOTE: The setting can be at a higher level if shredding tissue is more fibrous.
2.7.7. Run the shredder driver for approximately 10–12 s by pressing the trigger of the shredder driver, while maintaining the setting at 2.
2.7.8. Look at the tube to check whether all or most of the solid mass of the tissue is shredded through the lysis disk and passed into the upper chamber of the shredder chamber by looking at the tube. NOTE: It may be necessary to shred the tissue sample for a longer period. Simply return the tube to the shredder holder and continue the process. Most kinds of tissue samples require only 10–12 s of shredding. While all shredding will produce some heat due to friction, this short processing time will not significantly heat the sample.
2.7.9. After shredding the sample, lift the shredder chamber out of the holder unit. Use the tube tool to remove the shredder cap and place it on a clean surface for later use. NOTE: The shredder cap should be considered contaminated with sample extract, and depending on the sample type, may be contaminated with infectious materials. Do not use the same shredder cap for any other tissue samples to prevent cross-contamination.
2.8. Transfer the shredded heart tissue into the third beaker with 40 mL of washing buffer
and stir bar and stir the tissue for 5 min at a medium speed while on the stir plate.
2.9. Filter the suspension through a nylon mesh monofilament (74 μm pore size) and discard the filtrate. Wash the shredded tissue on the filter 3x with 10 mL of washing buffer. NOTE: Make sure that the nylon mesh monofilament is pre-wetted with water.
2.10. Transfer the washed and shredded tissue into a 50 mL beaker with 20 mL of washing buffer and place the beaker in the ice bath on the magnetic stirrer. Add 0.5 mL of trypsin solution while constantly stirring for 15 min. Approximately halfway through the incubation (at about 7.5 min), transfer the tissue suspension into a loosely fitted handheld glass-on-glass homogenizer that will hold 0–15 mL of volume. **(Figure 1B)**
2.11. Gently homogenize with 4–5 strokes to disperse the suspension without producing froth. Place the homogenate into a 50 mL beaker and gently mix on a stirring plate for 15 min at 4 °C.
2.12. After 15 min of incubation, add 10 mL of the isolation buffer that contains the trypsin inhibitor into the beaker containing the homogenate and incubate for 1 min at 4 °C while gently stirring.
2.13. Filter the suspension again through a nylon filter and save the filtrate in a 50 mL conical tube.
2.14. Collect the particulate fraction left on the nylon filter and place it in the glass-on-glass homogenizer. Add 15–20 mL of the washing buffer and gently homogenize by hand for 25–30 s. NOTE: The filtrate flows very slowly at this step. To increase the filtrate flow, place the funnel tip touching the inside of the conical tube.
2.15. Combine the homogenate with the filtrate, balance the tubes, and centrifuge at 600 x *g* for 15 min at 4 °C.
2.16. Filter the resulting supernatant into a new conical tube using a nylon filter to avoid contamination by the pellet. Run the centrifuge again at 8,500 x *g* for 15 min at 4 °C. NOTE: Use a plastic transfer pipette to remove the supernatant and sacrifice 0.2 mL of supernatant from the top of the fluffy pellet.
2.17. Discard the supernatant and rinse the pellet 2–3 times with 1 mL of the isolation buffer. Ensure to discard the fluffy white outer rim layer each time. NOTE: Wash the pellet by adding the isolation buffer using a pipette with 1 mL tip, or a new transfer pipette. Add the buffer to the side of the tube, not directly over the pellet, to prevent breaking the pellet and prevent the contamination of the supernatant. Next, aspirate the buffer with a transfer pipette and repeat the process two more times.
2.18. Add 5 mL of the isolation buffer to each tube and break up the pellet using a pipette with a 1 mL tip, or a new transfer pipette.
2.19. Dilute the suspension with 20 mL of additional isolation buffer and run the centrifuge again at 8,500 x *g* for 15 min at 4 °C.
2.20. Discard the supernatant and resuspend the pellet in 5 mL of the resuspending buffer. Then, add 15 mL more of the buffer for a total of 20 mL, and centrifuge at 8,500 x *g* for 15 min. at 4 °C. Discard the supernatant. The resulting brown pellet contains washed intact mitochondria.
2.21. Resuspend the mitochondria-enriched pellet in 250 μL of the resuspending buffer, including protease inhibitors. Aliquot in 25 μL, quick freeze in liquid nitrogen, and store in a −80 °C deep freezer for several years. NOTE: Spare 5 μL of mitochondria-enriched preparation for the protein analysis assay. For a BN-PAGE (Blue native-PAGE) study, aliquot the resuspended pellet in 50 μg of mitochondrial protein per tube. Centrifuge at 8,500 x *g* for 15 min at 4 °C. Carefully aspirate the supernatant and store the mitochondrial pellet (50 μg protein/aliquot) in a −80 °C deep freezer for several years.

**Figure 1:**
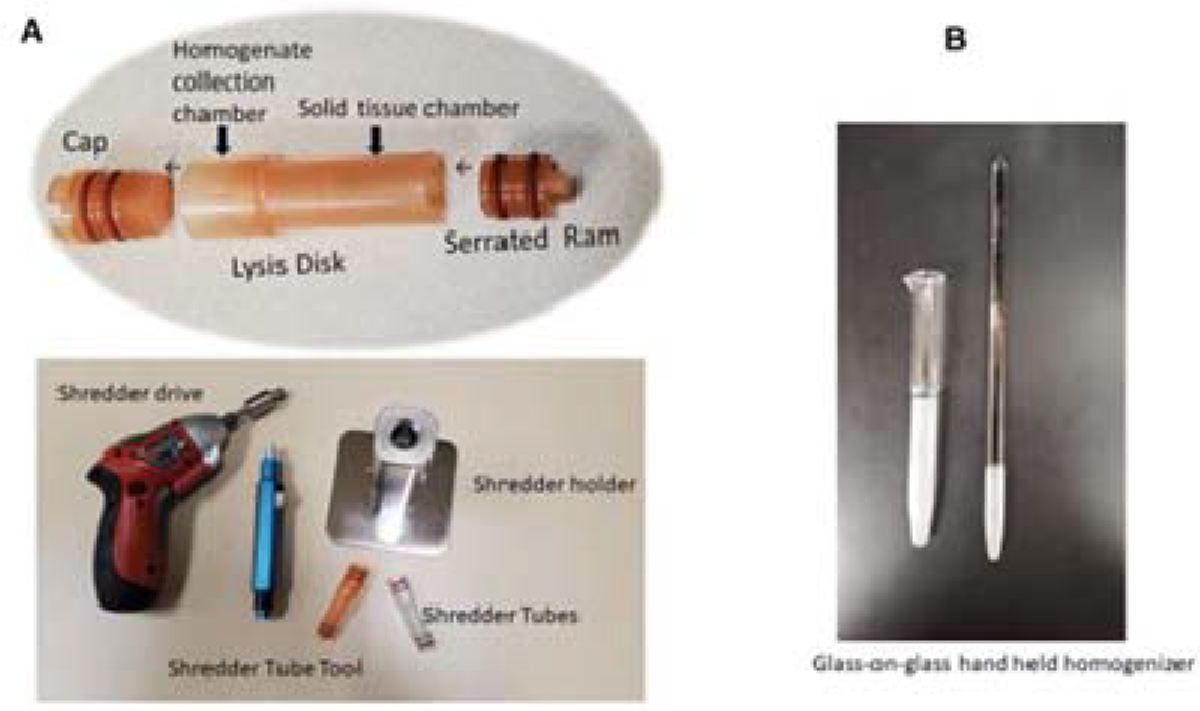
Tissue shredder chamber (tube) for use with the shredder. (**A**). Tissue homogenization in the shredder tube requires about 10–12 s per tissue sample. Tissue homogenization is completed once the homogenized tissue passes through the perforated disk into the upper chamber. (**B**. A glass-on glass hand-held homogenizer

### 3. Mitochondrial electron transfer complex (ETC) assessment

NOTE: Individual mitochondrial ETC proteins (i.e., Complex-I, II, III, IV, V) can be assessed by standard immunoblotting assay. Due to the variety of samples (four time points: 48 h, 3 months,6 months, 9 months), two experimental conditions (exercised and sedentary), and a number of immunoprobing antibodies (five antibodies), a multiplexed immunoblotting is recommended. Perform multiplexed immunoblotting as follows.

3.1. Resolve 50 μg of mitochondria enriched protein sample/well in 4%–20% gradient polyacrylamide gel electrophoresis (100 V, 90 min).
3.2. Transfer the proteins onto a PVDF membrane (75 V; 60 min)
3.3. Block the membrane in a blocking buffer solution for a minimum of 1 h at 4 °C. NOTE: Blocking time can be extended for up to overnight at 4 °C. The blocking buffer **(Table of Materials)** contains non-mammalian ingredients (i.e., fish proteins) that are less likely to have specific binding interactions with antibodies as well as other mammalian proteins present in typical methods.
3.4. Probe the membrane with the antibody cocktail including anti-Complex-I, II, III, IV, V antibodies and incubate overnight on an orbital shaker at 4 °C.
3.5. Next day, wash the membrane and probe with IR-tagged secondary antibody for 2 h on an orbital shaker at RT (room temperature).
3.6. Wash five times with 1x TBS-T (TBS containing Tween 20), and once with 1x TBS (Tris buffered saline). NOTE: For the last wash, do not include detergent in the TBS buffer.
3.7. Image the blot in an Image analyzer.

### 4. The mitochondria supercomplex assessment by Blue Native Poly Acrylamide Gel Electrophoresis (BN-PAGE)

NOTE: A supercomplex profile of ETC can also be assessed by employing the BN-PAGE technique.

4.1. Prepare the sample buffer cocktail for 20 μL per 50 μg mitochondria-enriched pellet protein (7 μL of water + 5 μL of 4x native polyacrylamide gel electrophoresis sample loading buffer + 8 μL of 5% Digitonin + 2 μL of Coomassie G-250). NOTE: Make a stock sample buffer cocktail depending on the number of samples available (50 μg protein/sample). Choose appropriate digitonin amounts based on the protein concentration of the mitochondria-enriched pellet (**Table 1**)^8^.
4.2. Heat the digitonin stock at 95 °C for 3 min, mix, and cool on ice before making the master mix.
4.3. Add the Coomassie G-250 sample additive just before loading the samples into the gel.
4.4. Add 20 μL of the sample buffer cocktail in a tube that contains 50 μg of mitochondrial protein pellet. Solubilize/resuspend the pellet by gently mixing the pellet with a pipette without generating froth.
4.5. Incubate the mixture on ice for 20 min to achieve maximum solubilization of the mitochondria-enriched pellet. From time to time, gently flick the tubes to enhance solubilization.
4.6. While waiting for incubation, prepare the native PAGE anode buffer/1x running buffer and the dark blue colored cathode buffer.
4.7. Centrifuge the solubilized mitochondria-enriched pellet at 20,000 x *g* for 10 min at 4 °C.
4.8. After centrifugation, collect 15 μL of the supernatant into new tubes placed on ice.
4.9. Add the appropriate amount of 5% Coomassie G-250 reagent to the supernatant obtained in step 4.8 (refer to **Table 1**). NOTE: The 5% Coomassie G-250 volume should be such that the final concentration of the dye is ¼ the detergent concentration. Perform this step just before loading the samples into the gel.
4.10. Pour a 3%–8% gradient gel, remove the comb, and wash out the wells of the gel with 1x running buffer three times. NOTE: The gel best polymerizes if poured a day before.
4.11. Set up the electrophoresis system and place the gel in the apparatus.
4.12. Fill the sample wells with the dark blue colored cathode buffer.
4.13. Load 10 μL of the unstained protein standard as a native molecular weight marker on the first and last well of the gel.
4.14. Load the entire amount of the sample supernatant + Coomassie sample additive prepared in step 4.9 above into the wells using a flat-head gel-loading tip.
4.15. Carefully fill the mini cell buffer dam with approximately 60 mL of dark blue colored cathode buffer without disturbing the samples in the wells, and then fill the buffer tank with the native PAGE 1x running buffer.
4.16. Turn on the power supply and electrophorese the samples at 150 V for approximately 30 min.
4.17. Remove the dark blue color cathode buffer from the buffer dam (inner chamber) with a transfer pipette or aspirate the sample wells. Fill the inner chamber with 150–200 mL of light blue colored cathode buffer and run the electrophoresis at 250 V for 60 min. NOTE: The electrophoresis time may be determined by checking when the dye front migrates just 0.5 cm above the bottom of the gel cassette.
4.18. Remove the gel from the cassette and place it in a clean plastic container. Wash the gel with good quality distilled water for 15 min on a rocker/orbital shaker. NOTE: To remove the gradient gel from the cassette, handle the cassette from the bottom, which is stronger, to minimize the chances of breakage.
4.19. Pour off water, immerse the gel in a sufficient amount of Coomassie GelCode Blue stain reagent, and incubate for 1 h on an orbital/rocker at room temperature. NOTE: Mix the Blue stain reagent **(Table of Materials)** solution before use by gently swirling the bottle several times.
4.20. Decant the dye solution, add distilled water, rinse several times with water, and then put on the orbital/rocker for destaining. NOTE: Insert a piece of sponge (~3 cm x 0.5 cm) into the gel destaining container and leave the gel for overnight destaining. The sponge adsorbs the dye particles without requiring multiple water changes.
4.21. Analyze the gel in an image analyzer.

**Table 1:** Sample Buffer Cocktail Preparation with two different detergent/protein ratio. To achieve maximum solubilization of membrane proteins from the mitochondrial suspension, the digitonin/mitochondrial suspension protein concentration should be adjusted to between 4–8 g digitonin/g of protein. In the heart tissue, 400 μg digitonin for 50 μg of mitochondrial suspension proteins (i.e., 8 g digitonin/g of mitochondria protein) were used to maximize the solubility of the supercomplexes from the mitochondria suspension.

| Table-1 : Sample Buffer Cocktail Preparation with two different detergent/protein ratio |  |  |  |  |  |  |  |  |  |  |
| --- | --- | --- | --- | --- | --- | --- | --- | --- | --- | --- |
| Protein | Digitonin/protein ratio (g/g) | | 4 X Sample Buffer $\mu\text{L}$ | | 5% Digitonin ($\mu\text{L}$) | | Water ($\mu\text{L}$) | Final volume ($\mu\text{L}$) | | 5% Coomass sample addit |
| 50 $\mu\text{g}$ | 8 | | 5 | | 8 | | 7 | 20 | | 2 |
| 50 $\mu\text{g}$ | 4 | | 5 | | 4 | | 11 | 20 | | 1 |

**Table 2:**
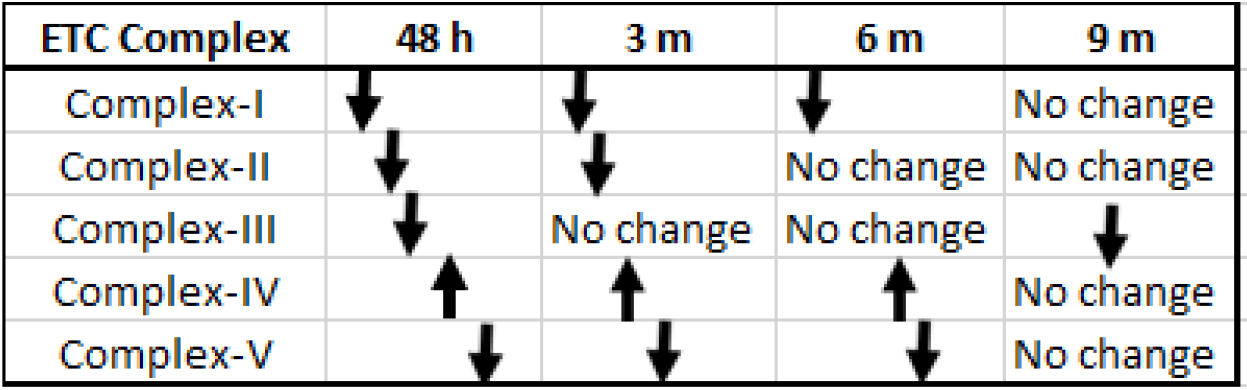
Mitochondrial electron transport chain protein complexes profile. The table summarizes the profile of mitochondrial ETC complex levels in the exercised group of sows. All complexes, except Complex-IV, showed a decreasing pattern across the age groups.

### 5. Mitochondria supercomplex assessment by immunoblot blot analysis

5.1. Perform a new set of BN-PAGE without staining the gel at the end of the electrophoresis.
5.2. Follow the same procedure as in section 3 (mitochondrial electron transfer complex (ETC) assessment) to complete the immunoblotting.

## REPRESENTATIVE RESULTS

Following the protocol, a good yield of mitochondria-enriched protein mixture from heart tissue was prepared. Approximately 15 mg/mL of mitochondria-enriched protein mixture was obtained from an average of 1.2 g frozen heart tissue harvested from the offspring of the sow. Observations indicated that less than 0.5 g of frozen heart tissue did not yield a sufficient amount of mitochondrial-enriched protein mixture to carry out a BN-PAGE assay. The amount of mitochondrial preparation was sufficient to perform (i) a standard immunoblot analysis for assessing the individual electron transfer complexes (ETC) (i.e., Complex-I, II, III, IV, and Complex-V), (ii) a mitochondrial supercomplex assessment by BN-PAGE and immunoblot analysis, and (iii) limited enzymatic assays for the select complex. The cocktail of antibodies raised against individual ETC protein complexes provides technical feasibility that each antibody will recognize its own target protein in one single immunoblot assay.

Mitochondria-enriched protein was prepared from the left ventricle of the heart, obtained from the offspring of the sow. The offspring were born to two groups of sows: (1) one group was exercised during pregnancy and (2) one group was sedentary. The heart tissues of both the groups were harvested at different time points after the birth (48 h, 3 months, 6 months, and 9 months). Whether maternal exercise has a positive impact on mitochondrial ETC of the offspring during gestation was the hypothesis that was tested in this study. Four heart tissues were comfortably processed on any given day due to the lengthy mitochondria-enriched isolation process.

**Figure 2** represents a typical ETC protein profile. Antibody cocktails allowed for immunoblotting in a multiplex format. Each antibody recognized its own target protein, providing an acceptable resolution. The protein band intensities were digitally analyzed. Tissues obtained from the offspring of exercised sow presented relatively reduced levels in some of the ETC complexes **(Supplemental Figures 1A–E).** The mitochondria-enriched preparation must be resolved in a gradient gel (4%–20%) as the immunoprobing was performed in a multiplexing format. This approach will eliminate the necessity of running several fixed percent of SDS/PAGE gels and run-to-run variability.

**Figure 2:**
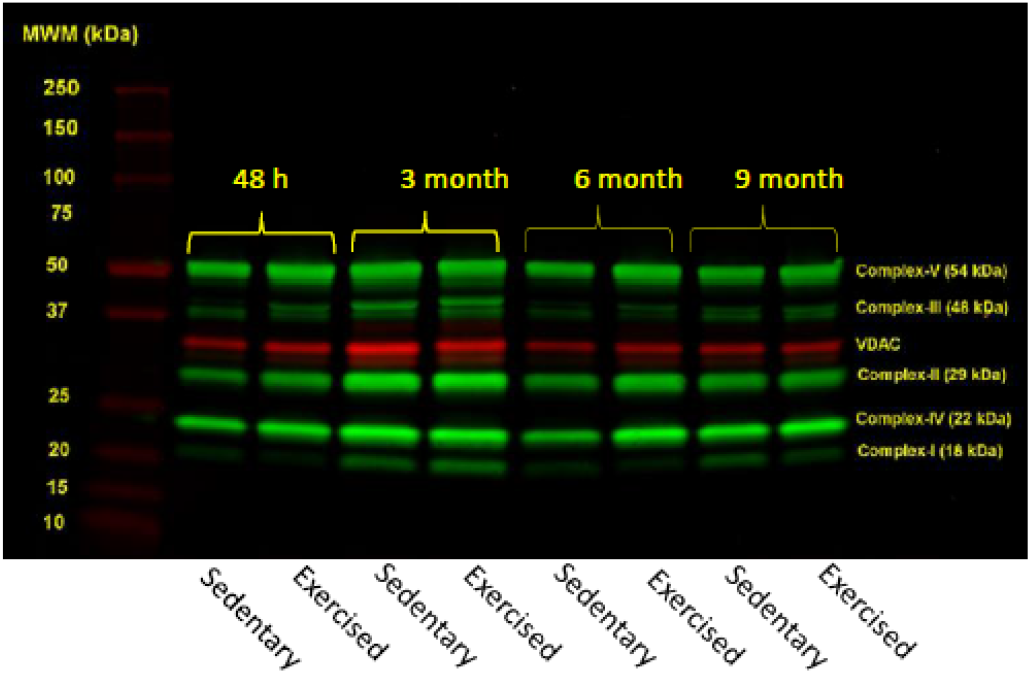
Immunoblot profile of mitochondrial ETC complexes from offspring heart tissue. The mitochondrial-enriched protein mixture was resolved in a 4%–20% gradient gel under reducing conditions. Proteins were immunoprobed by a commercial antibody cocktail, and anti-VDAC antibody was used for loading control (red color band). The cocktail of antibodies was raised against bovine heart Complex-I (antigen NDUFB8), bovine heart Complex-II (antigen C-II30 (FeS), bovine heart Complex II, III (antigen C-III-Core 2), human Complex IV, subunit II (antigen C-IV-II), and bovine heart Complex-V (antigen C-V-α). The secondary antibody was labeled with an infrared tag, and the image was analyzed using an image analyzing software. Individual Complex profiles across the age groups are provided in **Supplemental Figure 1 (A–E)**.

Mitochondrial proteins were resolved in a 3%–8% gradient BN-PAGE **(Figure 3A)** and immunoprobed with antibody cocktails. In both exercised and sedentary groups, multiple supercomplex formation was observed **(Figure 3B).**

**Figure 3:**
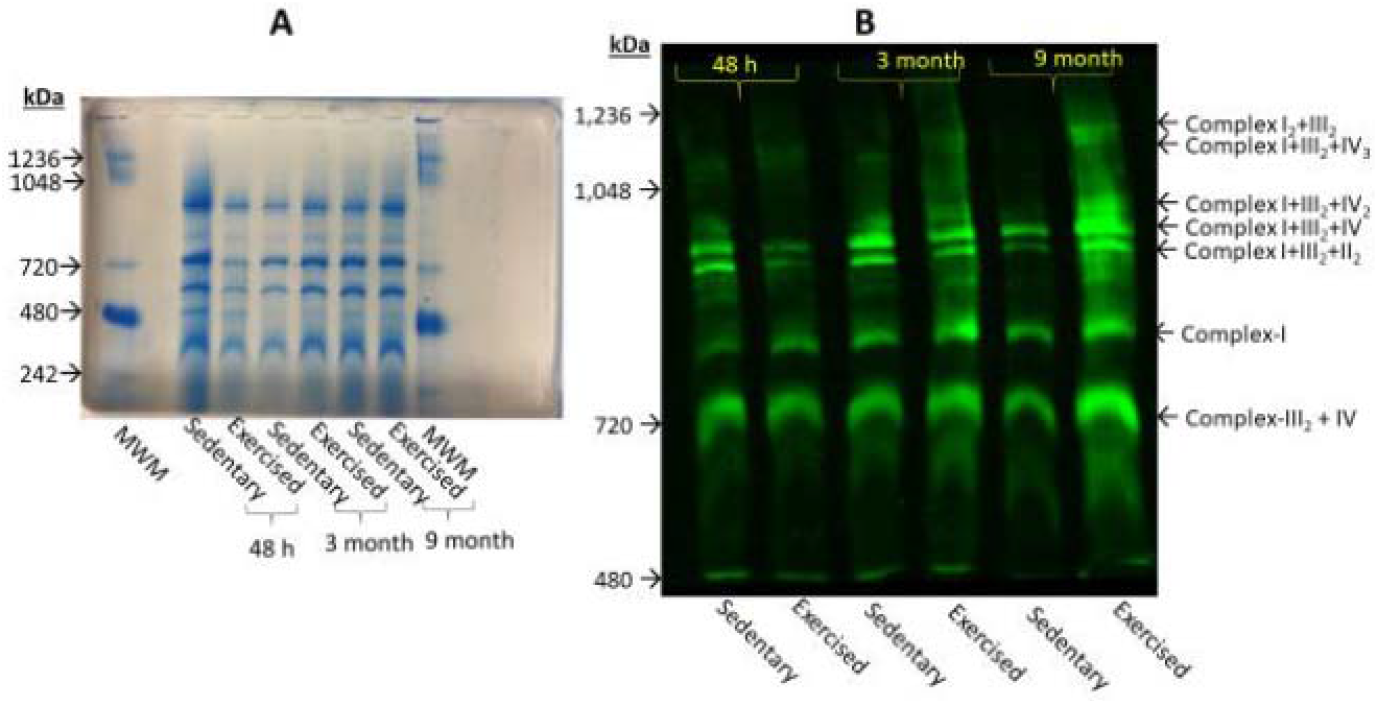
Mitochondrial supercomplex profile by Blue-Native Polyacrylamide gel electrophoresis (BN-PAGE). **(A).** Mitochondrial-enriched protein mixtures were resolved in 3%–8% BN-PAGE, and stained with a Coomassie blue reagent (Gel Code, Blue Stain Reagent). (B). Mitochondrial-enriched protein mixtures were resolved in a 3%–8% BN-PAGE, transferred onto a PVDF membrane, and immunoprobed with the antibody cocktails for supercomplexes. In both assays, the protein load was 150 μg/well. A 6-month data collection point was not included due to an insufficient protein yield during the preparation. The most prominent supercomplex increase observed was at 9 months in the exercised group compared to the sedentary group.

A different gradient gel format (3%–8%) was prepared due to the large size of the supercomplexes that can be resolved in BN-PAGE. Mitochondria-enriched preparation described in this protocol was used for measuring mitochondrial ETC complex activities. **Figure 4** represents data recorded for Complex I–II activity assessment. This assay was performed according to the established protocol^3^.

**Figure 4:**
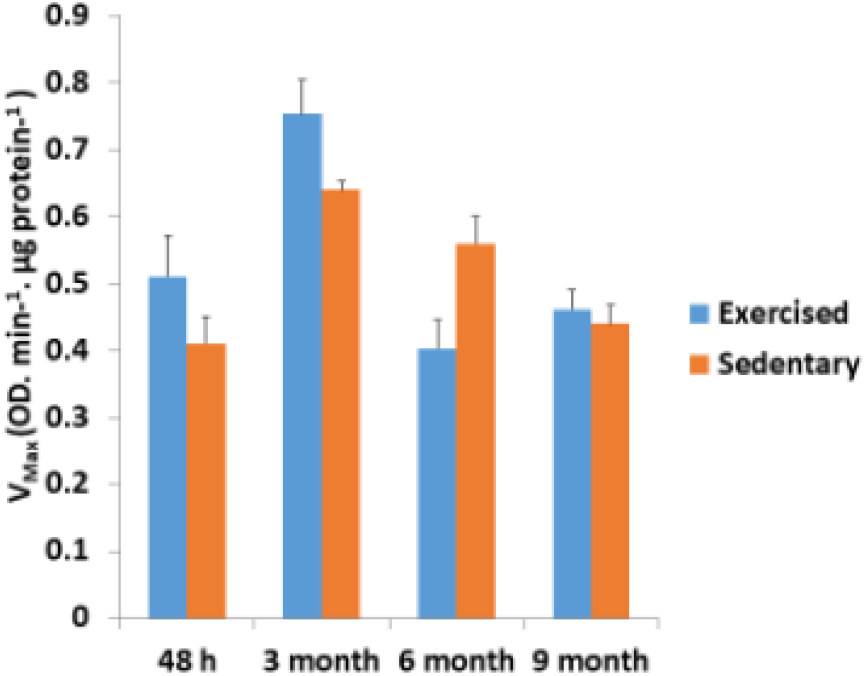
Complex I–II activity measurements. Mitochondria-enriched protein mixtures obtained from different age groups (48 h, 3 months, 6 months, and 9 months) were analyzed for the Complex I–II activity using a kinetic assay. The exercised group showed an increased Complex I–II activity as compare to that of the sedentary group. The difference between the two groups was not statistically significant according to a paired *t*-test (P > 0.05). The error bars depicted in the figure represent the standard error of the mean (SEM) for n = 4.

## Supporting information

Supplemantal Figure-1 (combined in one panel)

---

**Supplemental Figure** 1: **(A)** Complex-1 profile across age groups. In general, offspring born to the exercised sow presented with lower levels of Complex-I compared to that of the sedentary group (n = 4). This was more pronounced in the 9-month group. (B) Complex-II profile across age groups. All age groups showed low levels of Complex-II in the exercised group with the exception of three months cohort (n = 4). **(C)** Complex-III profile across age groups. Only the 9-month age group showed lower levels of Complex-III in the exercised group as compared to that of the sedentary group (n = 4). (D) Complex-IV profile across age groups. A slight increase in the Complex-IV protein levels of the exercised group was observed (n = 4). (E) The Complex-V (ATP synthase) profile across age groups. Both exercised and sedentary groups exhibited low levels of Complex-V except at the 48 h data point (n = 4).

## DISCUSSION

The critical steps for this protocol are indicated here. First, tissue homogenization should be carefully handled so that excessive sheer effects will not be applied during the tissue homogenization process. A tissue shredder should be used, which is a part of pressure cycling technology (PCT) for initial tissue homogenization^9^. This step will reduce the excessive stroke cycle of glass-on-glass homogenizer **(Figure 1B)** that may further destroy already fragile mitochondria due to frozen tissue conditions. Second, the frozen tissue weight will be important to obtain sufficient amounts of the mitochondria-enriched preparation. The recommended starting tissue amount will be greater than 0.8 g, depending on the source of the tissue. The heart tissue is very rich in mitochondria^10,11^; therefore, the recommended amount will be sufficient to obtain a mitochondria-enriched preparation. Third, all procedures should be performed in a cold room. If a cold room is not available, a large-sized ice bath will be sufficient.

The feasibility of obtaining mitochondria-enriched preparations from previously frozen heart tissue has been demonstrated in this study. This protocol will be essential for providing access to archived frozen tissue from which mitochondria can be isolated and from which mitochondrial ETC enzyme complexes can be assessed. A recent study demonstrated that mitochondrial respiration from previously frozen tissues and cells could be measurable as well^2^. Because of this advancement, investigators will be able to utilize previously collected biosamples for further analyses.

To assess the profile of individual ETC and supercomplexes is made easier by performing classical immunoblotting in a multiplexing format that utilizes multiple antibodies during the immunoblotting procedure on the same PVDH membrane. It is critical for the user to determine the amount of protein to be loaded into gels for SDS-PAGE and BN-PAGE. The resuspension volume should be as minimal as possible in order to obtain high protein concentrations so that the user will not have a problem with sample loading volume. The protein amount to be loaded in BN-PAGE depends on the yield of the mitochondria-enriched preparation. A 150 μg of mitochondria-enriched protein mixture was loaded per well in BN-PAGE due to the small amount of original heart tissue. The 50–150 μg mitochondria-enriched pellet preparation would be sufficient for 1 mm thick gel. The user may prefer a 1.5 mm thick gel for sample loading; however, the resolution of protein bands may not be of acceptable quality. The less mitochondrial protein resuspension per well was not tested because antibody cocktails used in immunoblot assays were predetermined in their concentration; however, users can make their own antibody cocktail to increase the signal/noise rate.

A fully polymerized gradient gel (3%–8%) for BN-PAGE gels is recommended. Gradient gels of 3%–8% have not been commercially available; therefore, homemade gradient (3%–8%) standard mini gels (9 × 6 cm) can be used. Pouring the gel 24 h before and allowing for the gel to polymerize overnight in RT will provide an acceptable gel formation. If a user needs better separation of supercomplexes, it is recommended to use larger gel formats (either midi gel 13.5 cm x 6.5 cm or long gel 28 cm x 16 cm).

The application of this protocol will allow investigators to isolate mitochondria and analyze the profile of ETC and supercomplexes in previously frozen archived tissues. All national and regional biospecimen depositories keep the biosamples in ultra-freezing conditions (i.e., −80 °C). Large number of tissues are being archived by institutions and drug companies for further studies and reagent sharing purposes. The National Institute of Health (NIH) mandates that any NIH-supported projects are required to share the reagents produced from the supported projects. These biorepositories are very valuable sources for obtaining frozen archived tissues for obtaining preliminary data for grant applications. To analyze enzyme activities of supercomplexes was not performed in this study; however, protocols are available to perform such assays^8^.

Using the methods described in this paper, individual ETC complexes and supercomplexes of mitochondria-enriched preparations obtained from the heart tissue of the offspring of sows were analyzed. Offspring born to exercised sows presented low levels of ETC during the first 6 months of their life and eventually leveling up by 9 months.

This observation may suggest that offspring born to the exercised sows during the pregnancy indicated fewer but more efficient ETC complexes in their heart tissue mitochondria **(Figure 4).** However, supercomplex formation in 9 months exercised group has been found to be elevated. Exercise may somehow affect the rearrangement of supercomplexes in conditions of increased energy demand^12^. In addition, Complex-V (ATP synthase) has exhibited a low profile in the first 6 months after birth. This response may be due to the probability of epigenetically modified ETC protein complexes. The small sample size (n = 4) for this study was not sufficient to arrive at such a conclusion; however, patterns of low ETC complex protein levels, high Complex I–II activity, and elevated supercomplex formation may suggest that bioenergetics in heart tissue of the progeny of exercised sows are influenced by exercise. Fewer but more efficiently working mitochondrial ETC complexes and the formation of increased supercomplexes are interesting phenomena to be investigated. This project is still in progress.

In summary, this protocol provides simple and straightforward mitochondria-enriched sample preparation from previously frozen archived heart tissue. Mitochondrial ETC complexes can be analyzed by standard immunoblotting with multiplexed antibody cocktails. BN-PAGE followed by immunoblotting can analyze the presence of supercomplexes. This preparation can also allow users to assess mitochondrial ETC complex activities.

## ACKNOWLEDGMENTS

This work was financially supported by Kansas City University’s intramural grant for Abdulbaki Agbas and Summer Research Fellowship for Daniel Barrera. The authors are thankful for Dr. Jan Talley’s editorial work.

