## Supplementary figures and images for "Measuring Mitochondrial Electron Transfer Complexes in Previously Frozen Cardiac Tissue from the Offspring of Sow: A Model to Assess Exercise-Induced Mitochondrial Bioenergetics Changes"

### Supplemantal Figure-1 (combined in one panel)

Supplemental Figure-1

**a**

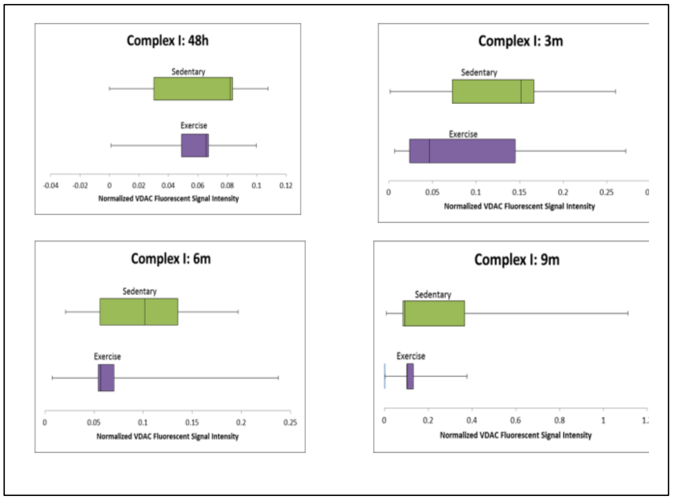


**b**

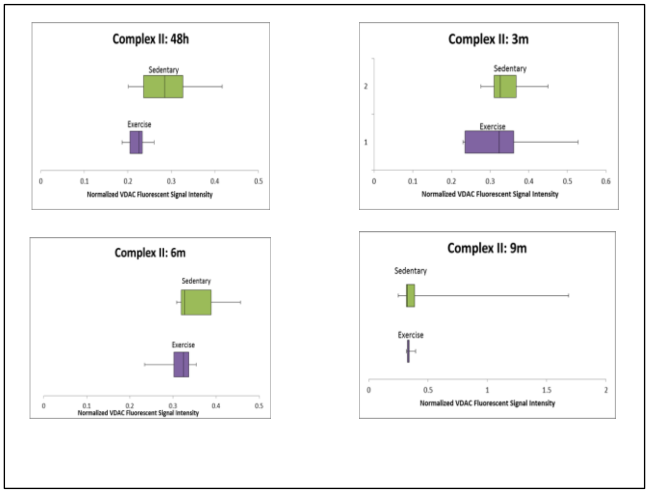


**c**

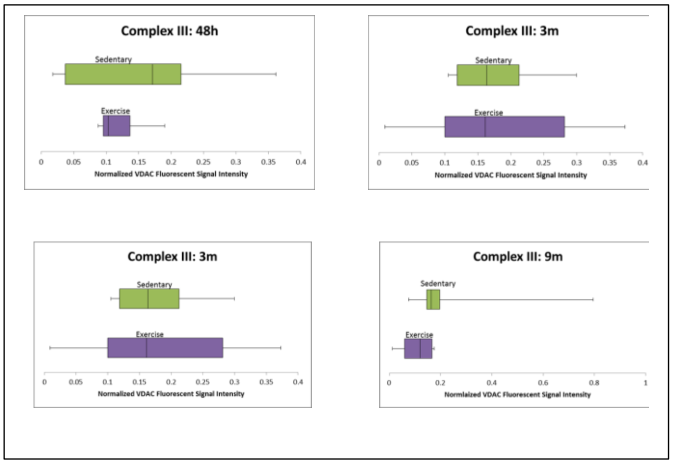


**d**

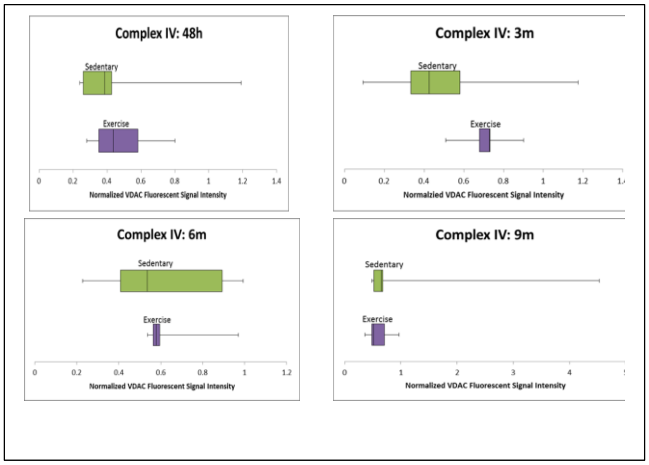


**e**

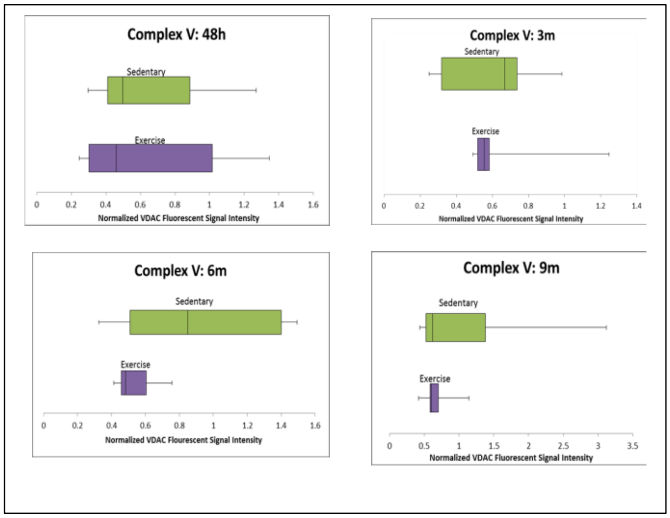
